# Structural Heterogeneity of Proteoform-Ligand Complexes in AMP-Activated Protein Kinase Uncovered by Integrated Top-Down Mass Spectrometry

**DOI:** 10.1101/2025.04.15.648965

**Authors:** Hsin-Ju Chan, Boris Krichel, Liam J. Bandura, Emily A. Chapman, Holden T. Rogers, Matthew S. Fischer, David S. Roberts, Zhan Gao, Man-Di Wang, Jingshing Wu, Charlotte Uetrecht, Song Jin, Ying Ge

## Abstract

AMP-activated protein kinase (AMPK) is a heterotrimeric complex (αβγ) that serves as a master regulator of cellular metabolism, making it a prominent drug target for various diseases. Post-translational modifications (PTMs) and ligand binding significantly affect the activity and function of AMPK. However, the dynamic interplay of PTMs, non-covalent interactions, and higher-order structures of the kinase complex remains poorly understood. Herein, we report the structural heterogeneity of the AMPK complex arising from ligand binding and proteoforms—protein products derived from PTMs, alternative splicing, and genetic variants—using integrated native and denatured top-down mass spectrometry (TDMS). The fully intact AMPK heterotrimeric complex exhibits heterogeneity due to phosphorylation and multiple adenosine monophosphate (AMP) binding states. Native TDMS delineates the subunit composition, AMP binding stoichiometry, and higher-order structure of AMPK complex, whilst denatured TDMS comprehensively characterizes the proteoforms and localizes the phosphorylation site. This is the first study to structurally characterize AMPK proteoform-ligand complexes. Notably, by integrating native TDMS and AlphaFold, we elucidate a flexibly connected regulatory region of AMPK β subunit that has been difficult to visualize with traditional structural biology tools. Our findings uncover previously unresolvable structural features of AMPK, offer new perspectives on protein kinase regulation, and establish a versatile framework for comprehensive characterization of proteoform-ligand complexes.

## Introduction

Kinases constitute one of the largest enzymatic superfamilies in eukaryotic cells and play key roles in cellular signaling by catalyzing reversible phosphorylation reactions^1–3^. AMP-activated protein kinase (AMPK), a heterotrimeric complex composed of a catalytic α subunit and two regulatory subunits, β and γ, serves as a master regulator of cellular energy metabolism^4–9^. Dysregulation of AMPK has been linked to various human diseases, including diabetes, cancer, and cardiovascular diseases; therefore, AMPK is a prominent therapeutic target^9–12^. The activity and function of AMPK are tightly regulated by post-translational modifications (PTMs) such as phosphorylation^13^, and by multiple non-covalent ligand-binding events including adenine nucleotide binding^14–20^. Understanding the dynamic interplay among PTMs, non-covalent interactors, and higher-order structures of the AMPK complex remains a significant challenge.

The limitations in conventional PTM detection and structural approaches have hindered the comprehensive characterization of AMPK complex. Western blotting^21,22^ and bottom-up proteomics^23,24^ have been used to detect PTMs in AMPK subunits but lack the capability to provide PTM information together with non-covalent interactors. X-ray crystallography and cryogenic electron microscopy (cryo-EM) have provided structural models of AMPK complex and their allosteric binding to small molecules^15–18^, but offer limited structural information for dynamic and flexible regions. For example, the carbohydrate-binding module (CBM) is flexibly linked to the rest of the complex, and its structure in non-activated AMPK heterotrimers remains unresolved^18,19^. Therefore, there is an urgent need for advanced technologies to uncover the dynamic and heterogeneous structures of AMPK, which is crucial for understanding its regulatory mechanisms and advancing therapeutic development.

Top-down mass spectrometry (TDMS) analyzes intact proteins without enzymatic digestion, enabling in-depth characterization of proteoforms—protein products arising from PTMs, alternative splicing, and genetic mutations^25^—and offering a bird’s-eye view of the proteoform landscape^26–32^. TDMS can directly determine the relative abundance of proteoforms, capture combinatorial PTMs, and localize PTM sites. Recently, native TDMS—which integrates native mass spectrometry (native MS)^33–36^ with traditional TDMS—has emerged as a powerful tool for characterizing proteoforms, non-covalent interactions, and higher-order structures of protein complexes, complementing other structural biology techniques^37–46^. In native TDMS, protein complexes are ionized under native-like conditions, preserving subunit-subunit and protein-ligand interactions. Following ionization, subunits are first ejected (complex-up analysis) to determine complex composition, and protein backbones are subsequently fragmented (complex-down analysis) to obtain sequence information^37^. Alternatively, intact protein complexes can also undergo direct fragmentation without prior subunit ejection using techniques such as electron- or photon-based dissociation, thereby offering insights into higher-order structures^38,47–50^. Protein kinases, which are commonly heteromeric complexes composed of catalytic and regulatory subunits with multiple PTMs and ligands, remain largely unexplored by TDMS-based approaches. Thus, it is important to establish TDMS-based methods for comprehensive structural characterization of the proteoform-ligand complexes of protein kinases and provide a key route to advance our knowledge in kinase regulation.

Herein, we established an integrated native and denatured TDMS strategy to uncover the heterogeneity and dynamic structure of AMPK complex. AMPK exhibits multiple proteoform-ligand complexes arising from phosphorylation and adenosine monophosphate (AMP) binding. Our results determine the subunit composition of the heterotrimeric complex, AMP binding stoichiometry, and localize the phosphorylation site. Furthermore, we elucidate a flexibly connected regulatory region of AMPK β subunit, which has been unresolved by traditional structural biology techniques, using native TDMS and AlphaFold^51,52^. Overall, this study demonstrates that the integrated TDMS method provides structural information of heteromeric protein kinase complexes that are complementary to conventional methods by revealing their proteoforms, ligand binding, and higher-order structures.

## Results

### An Integrated TDMS Strategy for Structural Characterization of AMPK

To comprehensively characterize AMPK, we expressed an AMPK heterotrimeric complex and analyzed it using an integrated native and denatured TDMS platform. We first constructed the recombinant protein expression system based on the work published by Yan *et al.*^18^, where a tricistronic plasmid encoding the sequence of human AMPK α1, β2, and γ1 subunits was developed for cryo-EM structure determination. The α1 and γ1 isoforms are ubiquitously expressed across tissues, while β2 is the predominant isoform in skeletal and cardiac muscle^8,11^. SDS-PAGE analysis confirmed the expression of all three AMPK subunits and their purity (**Figure S1**). We leveraged the ultrahigh resolving power of a Fourier transform ion cyclotron resonance mass spectrometer (FTICR-MS) for comprehensive native and denatured TDMS analysis, while employing a quadrupole-time-of-flight mass spectrometer (Q-TOF) coupled with online reversed-phase liquid chromatography (RPLC) for rapid proteoform profiling and sequencing (**Figure 1a**). Native MS revealed the heterogeneity of AMPK proteoform-ligand complexes (**Figure 1b**). Furthermore, complex-up, complex-down, and native top-down analyses generated subunit dissociation and/or comprehensive MS/MS fragmentation to acquire primary-to-quaternary structural information (**Figure 1c**). Finally, denatured TDMS provided a holistic view of the proteoform landscape and enabled PTM site localization (**Figure 1d**).

**Figure 1.**
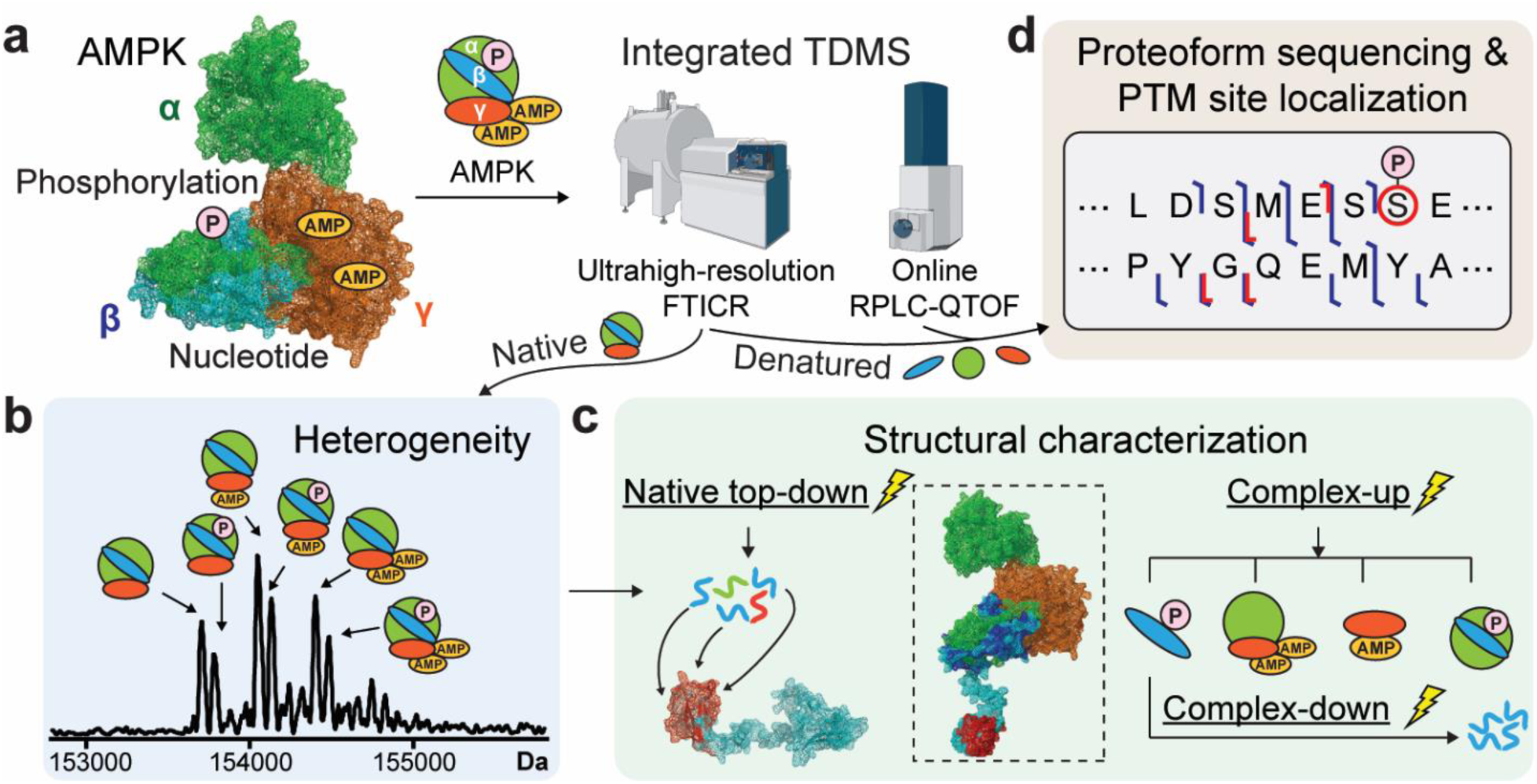
Characterization of the AMPK heterotrimeric complex by integrated top-down mass spectrometry (TDMS). (a) AMPK α1β2γ1 heterotrimer was characterized by integrated native and denatured TDMS. Native AMPK complex or denatured subunits were electrosprayed into mass spectrometers. An ultrahigh-resolution FTICR-MS was used for both native and denatured analysis, while a Q-TOF coupled with online RPLC was used for denatured subunit proteoform profiling. (b) The heterogeneous AMPK proteoform-ligand complexes were resolved by native TDMS. (c) Structural characterization of the AMPK complex using native top-down, complex-up, and complex-down analyses. (d) Proteoform characterization was enabled by denatured TDMS, providing sequence and PTM site localization information. PDB: 7M74, AlphaFold: O43741-F1-v4. P: phosphorylation, AMP: adenosine monophosphate.

### Heterogeneity of AMPK Proteoform-Ligand Complexes Resolved by Native MS

To elucidate AMPK proteoforms and ligand binding, the AMPK complex was buffer-exchanged into ammonium acetate and directly infused into an FTICR mass spectrometer. The native mass spectrum of the AMPK complex revealed a charge state distribution of 23+ to 29+ (5300 *m/z* to 6800 *m/z*) (**Figure 2a**), which corresponded to the ∼154 kDa AMPK αβγ heterotrimer (**Table S1**). The molecular mass confirmed that the complex was assembled as a 1:1:1 stoichiometric complex. The most abundant charge state 26+ of the AMPK heterotrimer was detected between 5910 *m/z* to 5950 *m/z*. Deconvolution of the native mass spectrum revealed the heterogeneity of the complex, which arose from six dominant AMPK proteoform-ligand complexes, each composed of a unique combination of PTMs and non-covalent interactors (**Figure 2b** and **Table S1**). Notably, the complex showed modification of up to one phosphorylation and binding of up to two AMP molecules. Next, we quantified individual proteoform-ligand complexes to determine the AMP binding and phosphorylation stoichiometry for the complex (**Figure 2c**). The relative proportion of AMP binding states for unbound, singly, and doubly bound complex species was 24%, 42%, and 33%, respectively. We also observed the complex was 59% unphosphorylated and 41% monophosphorylated. The comparison of the intensity ratios of the phosphorylated and unphosphorylated complex at different AMP binding states indicates that the AMP binding affinity is not affected by phosphorylation (**Figure S2**). Thus, native MS demonstrates that human AMPK exists as a highly heterogeneous protein assembly and exhibits diverse combinations of proteoform-ligand complexes.

**Figure 2.**
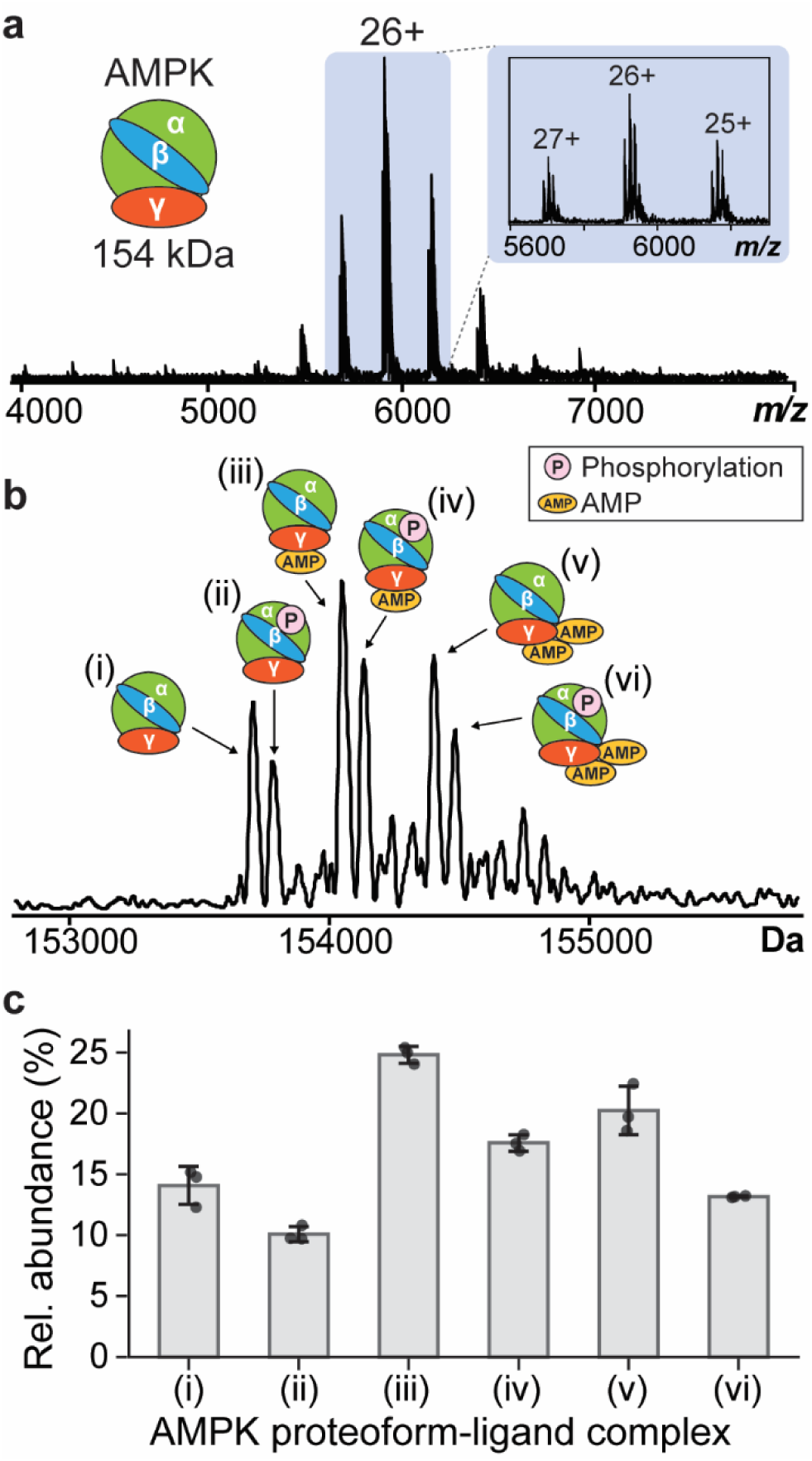
Native MS analysis using an FTICR-MS demonstrates the heterogeneity of AMPK heterotrimeric complex. (a) Native mass spectrum of AMPK αβγ complex. The inset shows a zoomed-in view of the three most abundant charge states *z* = 25-27+. (b) Deconvoluted native mass spectrum. Six peaks between 153 kDa and 155 kDa attributed to the AMPK complex are labeled with their corresponding proteoforms and bound ligands (P: phosphorylation, AMP: adenosine monophosphate). (c) Relative abundance of the six AMPK proteoform-ligand complexes. Data are presented as mean ± standard deviation (n = 3).

### Characterizing the Quaternary Structure of AMPK Complex via Native TDMS

We further performed complex-up analysis using collisionally activated dissociation (CAD) to elucidate the composition of the AMPK proteoform-ligand complexes (**Figure 3a**). The heterotrimeric complex dissociated into highly charged β and γ monomers, along with their charge-stripped αγ and αβ dimer counterparts (**Figure 3b** and **3c**). We then deconvoluted the mass spectrum from complex-up analysis and assigned each peak to its corresponding subunits (**Figure 3d-3g** and **Table S2**). Interestingly, phosphorylation was only observed on the β subunit (**Figure 3d** and **Figure S3a**). For the ejected γ monomer, the predominant proteoform observed was unmodified (**Figure 3e**). Notably, the AMP-bound γ was identified using increased instrument resolution (**Figure S3b** and **Table S3**). This finding confirms that AMP molecules bind to the γ subunit, consistent with previous studies^14,16,19^. We also identified low abundance of gluconoylation and phosphogluconoylation (**Figure S3b**), which are modifications commonly observed in overexpressed proteins and are often not considered biologically relevant^53^.

**Figure 3.**
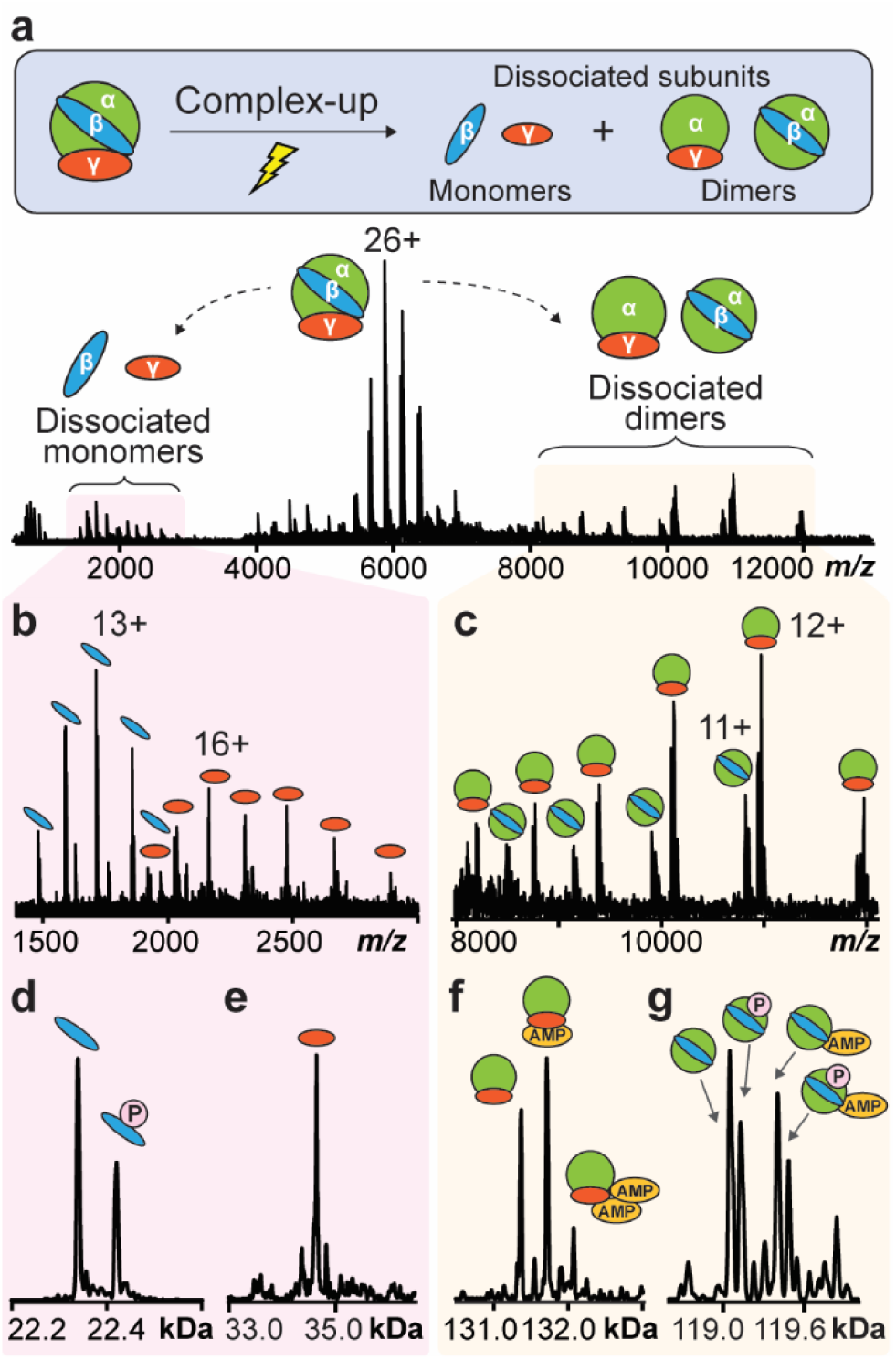
Complex-up analysis maps the PTMs and ligand binding to specific subunits. (a) Representative mass spectrum from complex-up analysis shows AMPK heterotrimer dissociated into monomers and heterodimers. Zoomed-in spectrum shows the charge state distribution of ejected (b) β and γ monomers as well as (c) αγ and αβ dimers. Deconvoluted mass spectrum of dissociated (d) β, (e) γ, (f) αγ, and (g) αβ subunits. P: phosphorylation, AMP: adenosine monophosphate.

Interestingly, we observed that AMP molecules were also bound to the αγ dimer (**Figure 3f**). Comparing the ejected γ and αγ subunits, the αγ subunit had a significantly higher AMP binding level than the γ subunit (**Figure 3f** and **Figure S3b**). The αγ dimer allowed binding up to two AMP molecules, with its binding profile consistent to that of the AMPK complex precursor. In contrast, the γ monomer bound to at most one AMP molecule, and the AMP-bound γ was present in low abundance. This difference in AMP binding cannot be attributed solely to the absence of the α subunit. Rather, it is a consequence of the CAD process, where ejected monomers like γ become highly charged CAD products that undergo partial unfolding, disrupting their ability to retain non-covalent interactions with ligands. Conversely, lower-charged CAD products are more likely to preserve their native-like structure and non-covalent interactions^54^. As a result, the charge-stripped αγ dimer maintains a higher AMP binding capacity. The observed differences in AMP binding reflect the structural integrity of the CAD products rather than an intrinsic requirement for the α subunit in AMP binding to γ. Moreover, the deconvoluted spectrum of the αβ subunit also confirmed that the phosphorylation was identified on the β subunit (**Figure 3g**). It should be noted that a small portion of αβ was bound to AMP molecules possibly due to rearrangements induced by collisional activation prior to the dissociation process^55–57^. Taken together, the unique dissociation profile of AMPK in complex-up analysis reveals subunit-specific PTMs, ligand binding, and subunit-subunit interaction. This analysis further provides insights into the distinct roles of each subunit in protein kinase function and regulation.

To gain additional sequence-specific information, we performed complex-down analysis of the dissociated AMPK subunits. The AMPK subunits were first dissociated using in-source CAD (**Figure S3**). We focused on isolating dissociated monomers at the lower *m/z* range (< 6000 *m/z*) for fragmentation due to the *m/z* limit of quadrupole isolation on our instrument. The ejected β (*z* = 12+) and γ (*z* = 16+) monomers were then isolated and fragmented using CAD in the collision cell (**Figure 4a, 4b, Figure S4** and **S5**). We identified 9 *b* and 4 *y* ions for the β subunit, and 10 *b* and 10 *y* ions for the γ subunit. All fragments were validated based on their mass accuracy and isotope distribution (**Figure 4c, 4d** and **Table S4**). Protein regions covered by identified fragments were illustrated on AMPK protein structures^18,51,52^ (**Figure 4e** and **4f**). For the β subunit, 5 more *b* ions were detected compared to *y* ions, suggesting the N-terminal domain may be slightly more prone to CAD fragmentation compared to the C-terminal domain. Meanwhile, backbone cleavages of the γ subunit were observed among all four cystathionine β-synthase (CBS) motifs, where mutations can occur and cause a variety of human hereditary diseases^15^. The results suggested that both termini of the γ monomer are easily accessed by CAD fragmentation. Fragmentation also localized methionine removal at the N-termini for both the β and γ subunits. Consequently, complex-up and complex-down analysis together provide unique insights into the primary-to-quaternary structure of AMPK heterotrimeric complex.

**Figure 4.**
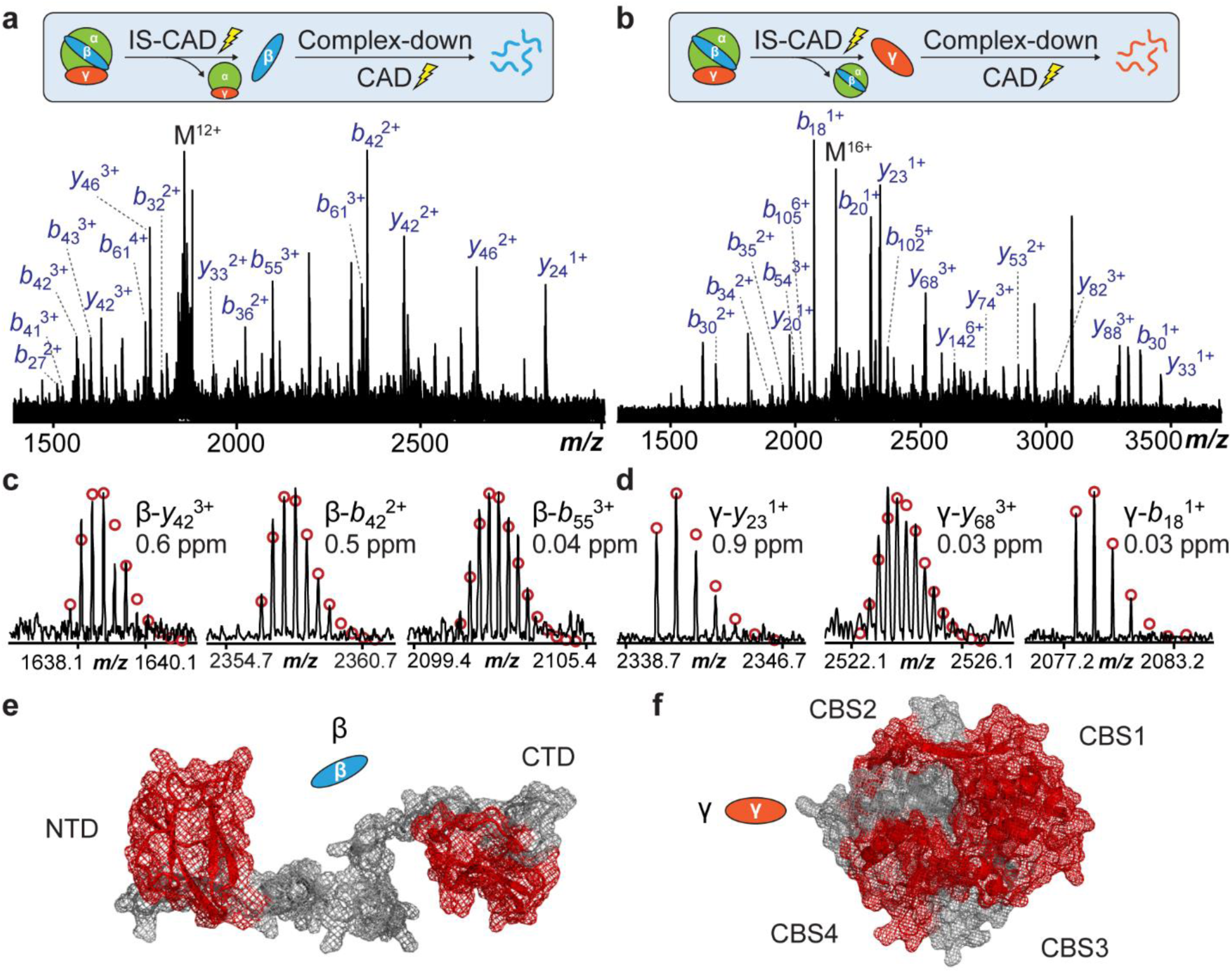
Complex-down analysis of ejected β and γ subunits provides sequence information. Complex-down mass spectra of the ejected (a) β (*z* = 12+) and (b) γ (*z* = 16+) subunit precursors. Representative fragment spectra of (c) β and (d) γ subunits. The isotopic fitting is shown in red circles and mass errors are reported. Structural representation of (e) β (AlphaFold: AF-O43741-F1-v4) and (f) γ (PDB: 7M74) labeled with fragments in complex-down analysis. The regions covered by identified fragments are labeled red. NTD: N-terminal domain; CTD: C-terminal domain; CBS: cystathionine beta-synthase motifs.

### Higher-Order Structural Elucidation of AMPK Complex Revealing the Dynamic Carbohydrate-Binding Module

Next, to further elucidate the higher-order structural characteristics of AMPK, we performed native TDMS using electron-capture dissociation (ECD) on the AMPK complex across all charge states (*z* = 23-29+). In the native top-down ECD spectrum, we detected the charge-reduced AMPK complex, with its three most intense charge states (*z* = 13-15+) appearing between 10000 and 12000 *m/z*, and ECD fragments observed at <2500 *m/z* (**Figure 5a**). Unlike the complex-down spectrum, no dissociated subunits were observed during native top-down ECD analysis. We detected fragments for all three subunits, including 14 *c* ions for the α subunit tagged with maltose binding protein, 45 *c* ions for the β subunit, and 1 *y* ion for the γ subunit (**Figure 5b**, **Figure S6** and **S7**). We mapped the fragment cleavage sites to a cryo-EM structure (PDB 7M74)^18^ of non-activated AMPK heterotrimer (i.e., not phosphorylated by upstream kinases) that was generated from the same construct used in our study. Interestingly, the cleavage sites on the β subunit could not be annotated in the resolved model, as they all occur in the N-terminal region, which is absent from the cryo-EM structure of our protein construct.

**Figure 5.**
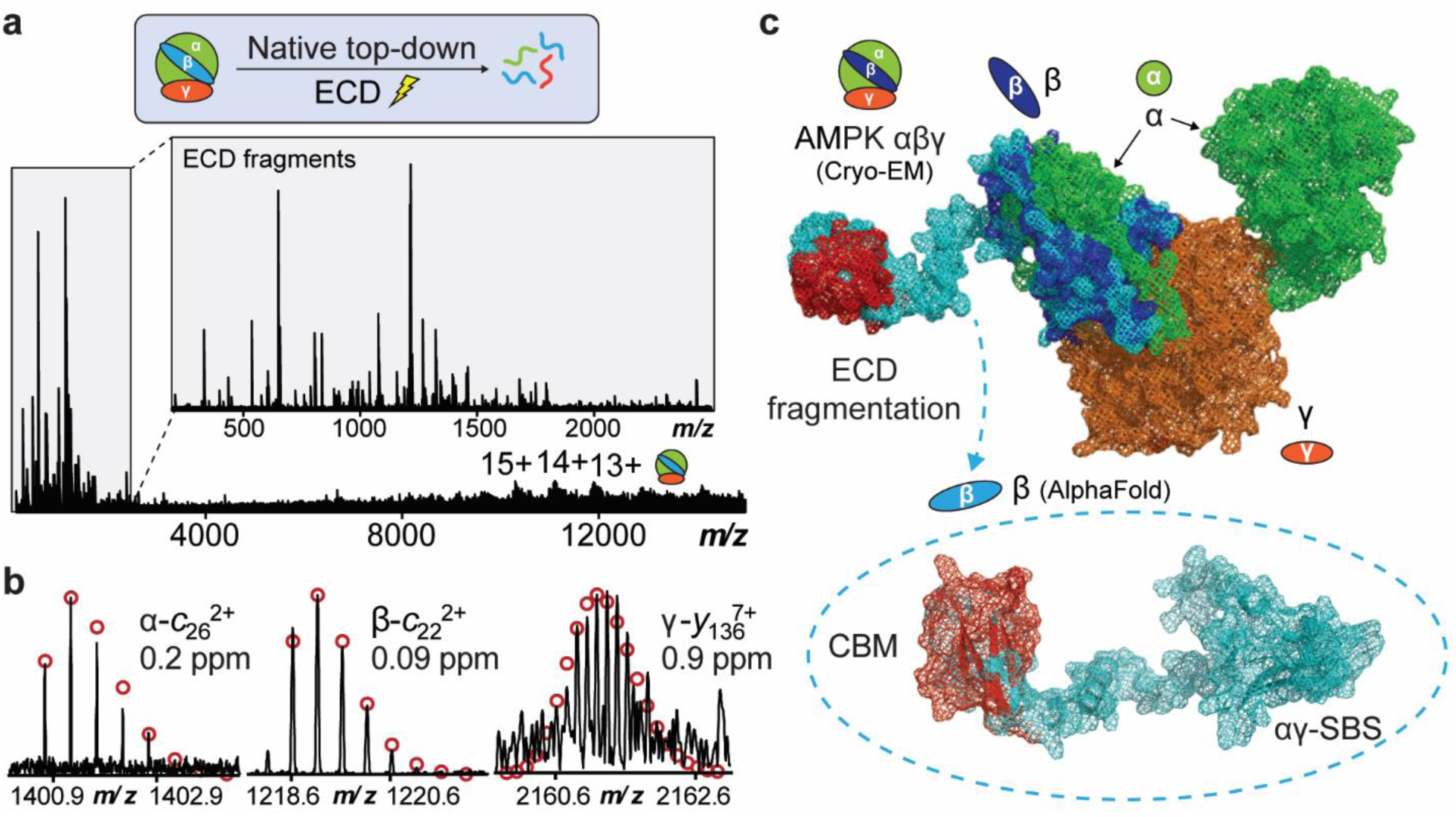
Native top-down ECD analysis uncovers a previously unresolved AMPK flexible region. (a) Representative mass spectrum of native top-down analysis using ECD shows AMPK heterotrimer charge-reduced species and ECD fragments. (b) Selected fragment spectra of three AMPK subunits. The isotopic fitting is shown in red circles and mass errors are reported. (c) AMPK structures annotated with ECD fragmentation sites. A chimeric model was constructed by aligning a cryo-EM structure of AMPK heterotrimeric complex (PDB: 7M74) and a predicted structure of full-length β subunit (AlphaFold: AF-O43741-F1-v4). Experimental and predicted structure of the β subunit is labeled in dark blue and cyan, respectively. Bond cleavage sites are labeled in red. CBM: carbohydrate-binding module, αγ-SBS: αγ subunit-binding sequence.

We therefore established a chimeric model composed of the cryo-EM structural model (PDB 7M74) and a full-length β subunit structure predicted by AlphaFold (AF-O43741-F1-v4)^51,52^ through aligning their β C-termini. We then mapped the ECD cleavage sites to this model (**Figure 5c** and **Figure S8**). In the resulting structure, the N-terminal region of the β subunit is solvent-exposed, consistent with our observation from the MS/MS fragmentation pattern. Notably, this region, known as the carbohydrate-binding module (CBM), has been reported to exhibit glycogen binding affinity and regulate the activity of AMPK^9,58–62^. Furthermore, it is also involved in forming the allosteric drug and metabolite (ADaM)-binding site, a binding pocket for pharmacological AMPK activators^63–65^. The CBM has been historically challenging to visualize using conventional structural biology tools, especially in AMPK complex not phosphorylated and activated by upstream kinases^18,19^. To the best of our knowledge, no structural characterization of the CBM in non-activated AMPK heterotrimers has been reported due to its high flexibility and solvent exposure. Here, we provide the first structural characterization of the CBM in a non-activated AMPK complex using native TDMS. In contrast, no cleavage sites were identified within the stable C-terminal region, which is known as αγ subunit-binding sequence (αγ-SBS) motif and tightly associates with the other two subunits. Altogether, these findings underscore the capability of native TDMS analysis in capturing dynamic structures and providing higher-order structural insights into the AMPK complex. Additionally, we establish a method for investigating the CBM, a key domain essential for AMPK regulation, with potential applications in studying its interactions with allosteric binders.

### In-Depth Characterization of AMPK Subunit Proteoforms through Denatured TDMS

Finally, we performed denatured TDMS to achieve in-depth characterization of individual subunit proteoforms. Previously, under native TDMS characterization, the α subunit did not dissociate as a monomer and therefore was not fully characterized. To characterize the α subunit, we first performed proteoform profiling using a Q-TOF mass spectrometer coupled with RPLC. Online RPLC produced baseline separation of the three subunits for MS detection (**Figure S9a**). The mass spectra were averaged over retention time windows and deconvoluted for proteoform identification (**Figure S9b-S9d)**. All proteoforms were identified with high mass accuracy. For the α subunit, the major proteoform (Expt’l: 96713.0 Da, Calc’d: 96713.6 Da) revealed the removal of the N-terminal methionine (**Figure S9b**). The α subunit was then fragmented using CAD, generating 16 fragment ions that confirmed the protein identity (**Figure S10**). As for the β and γ subunits, the β subunit proteoforms corresponded to N-terminal methionine removal and up to one phosphorylation, while the γ proteoforms coincided with N-terminal methionine removal and low-abundance of (phospho)gluconoylation, further validating what we observed in native TMDS analysis (**Figure S9c** and **S9d**).

To enable comprehensive proteoform characterization, we used ultrahigh-resolution FTICR-MS due to its exceptional resolving power for distinguishing overlapping ions. The stoichiometry of the unphosphorylated and monophosphorylated β proteoforms was 57% and 43%, respectively, similar to our native MS analysis (**Figure 6a** and **Figure S11**). Following intact MS analysis, we conducted tandem-MS (MS/MS) fragmentation using CAD and ECD for proteoform sequencing and PTM localization. For the β subunit, *b* ^12+^ was the first fragment counting from the N-terminus that contained phosphorylation (**Figure 6b**). The *b* ^12+^ ion in combination with *c*_98_ ^10+^, *z*_100_ ^9+^, and *y*_97_ ^10+^ demonstrated site-specific phosphorylation at Ser99 (Ser174 in canonical sequence), a previously reported Unc-51-like kinase 1 (Ulk1)-mediated phosphorylation site that potentially constitutes a negative regulatory feedback loop in autophagy^66^. Combining CAD and ECD spectra, we obtained 85% bond cleavage of the β subunit (**Figure 6c**). For the γ subunit, MS/MS fragmentation produced 53% bond cleavage (**Figure S12**). Overall, denatured TDMS provides a bird’s-eye view of the kinase proteoform landscape with high mass accuracy and enables proteoform sequencing as well as phosphorylation site localization.

**Figure 6.**
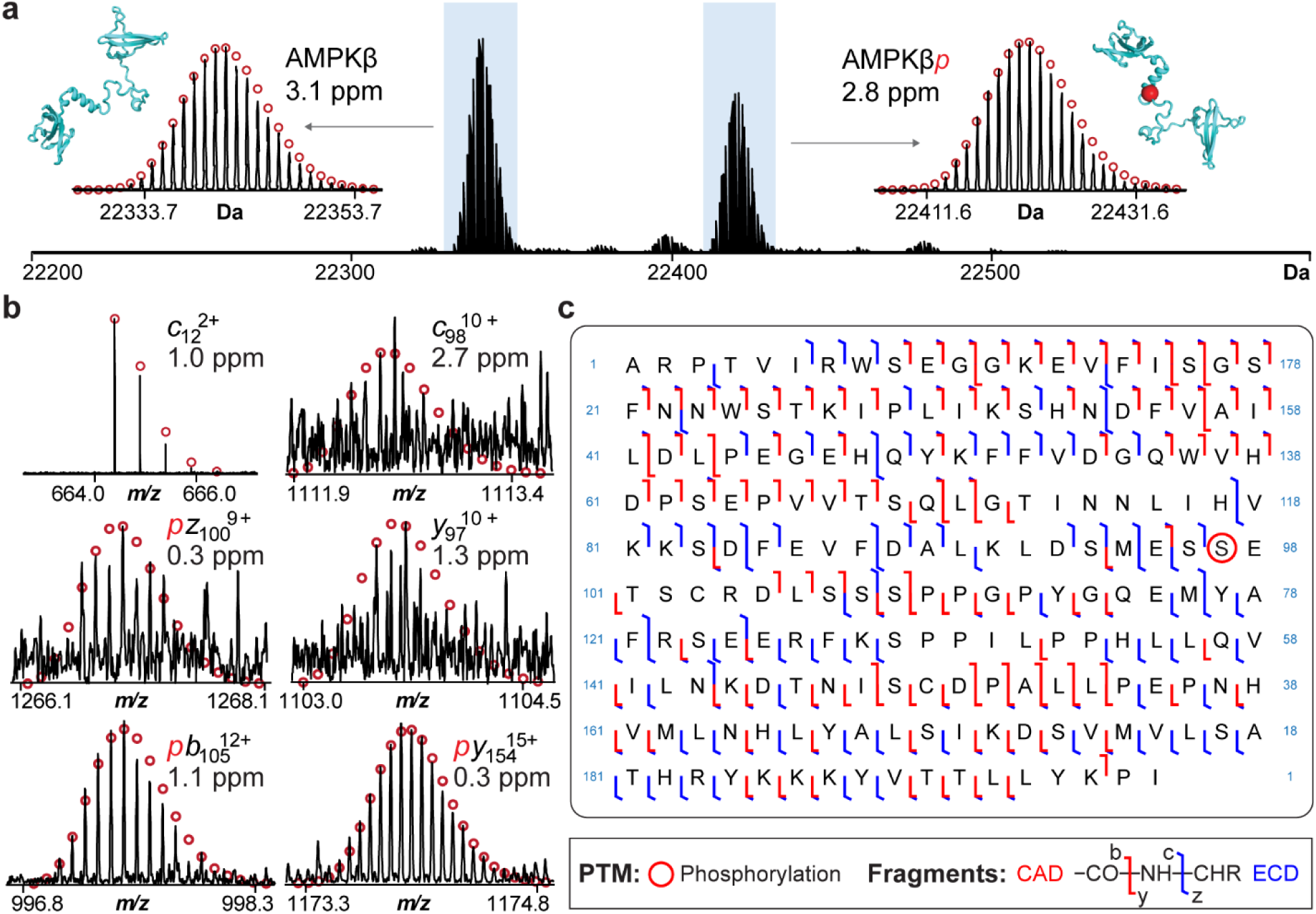
Denatured TDMS analysis localizes the phosphorylation site in AMPK β subunit. (a) Deconvoluted mass spectra of the AMPK β proteoforms acquired using ultrahigh-resolution FTICR-MS to achieve isotopic resolution. AMPKβ: unphosphorylated, AMPKβ*p*: monophosphorylated. Structural representation of the β subunit (AlphaFold: AF-O43741-F1-v4) with the experimentally defined phosphorylation site highlighted as a red sphere. The isotopic fitting is shown in red circles and mass errors are reported. (b) Representative CAD and ECD fragment spectra. (c) Sequence maps of the β subunit annotated with identified CAD and ECD fragments.

## Discussion

The integrated native and denatured TDMS platform reported here addresses key challenges in studying the interplay between kinase proteoforms, non-covalent interactors, and higher-order structures. For the first time, we demonstrate the TDMS-based structural characterization of heteromeric protein kinase complexes, revealing the heterogeneity and dynamic structure of AMPK proteoform-ligand complexes. This approach can provide complementary information to address the current knowledge gap in kinase biology.

Our study significantly advances the structural understanding of AMPK by providing insights previously unattainable with conventional structural biology tools^16–19^ and biochemical assays^15,20^. While these methods have characterized AMPK, they were unable to detect proteoforms and ligands within the complex simultaneously. In this study, we resolved six major proteoform-ligand complexes of AMPK and demonstrated that these complexes were composed of an unmodified α subunit, monophosphorylated/unphosphorylated β subunits, and unbound/singly or doubly AMP-bound γ subunits. Importantly, our results revealed the proteoform stoichiometry and the distribution of multiple ligand-binding states of AMPK complex.

Moreover, dynamic protein regions have remained challenging to study using conventional structural biology tools such as X-ray crystallography and cryo-EM^37,67^. According to available information on the Protein Data Bank, there has been no structural characterization of the CBM in non-activated AMPK heterotrimeric complex ever reported. Using native top-down ECD analysis assisted with AlphaFold, we successfully characterized CBM as a flexible and solvent-exposed region. Notably, the ECD fragmentation patterns of the β subunit in native and denatured TDMS analysis were significantly different, proving that the native-like structure of AMPK complex was retained in native TDMS. Thus, these data highlight the capability of native TDMS to elucidate the higher-order structures and offer a unique opportunity to study this dynamic region regulating AMPK activity. Given the significance of glycogen and allosteric activator binding in AMPK regulation, we also provide a potential approach for investigating their non-covalent interactions.

We previously characterized the catalytic domain of AMPK with C-terminal truncation using denatured TDMS and identified its phosphoproteoforms^68^. Nevertheless, the proteoform information of the other two regulatory subunits remained elusive. In this study, we achieved comprehensive characterization of all three AMPK subunit proteoforms expressed as a complex. We further localized the phosphorylation to the β-Ser174 in the canonical sequence, an Ulk1-mediated phosphorylation site for potential inhibition in autophagy^66^. Finally, we provided a strategy that directly establishes the relationship between kinase proteoforms and ligand binding by quantifying individual proteoform-ligand complexes. We found that the phosphorylation on β-Ser174 does not affect the AMP binding affinity to the complex.

In conclusion, the characterization of kinase complexes by the integrated native and denatured TDMS approach reveals new insights into kinase complex structures, proteoform heterogeneity, and ligand binding. By leveraging native TDMS, we successfully determine the non-covalent interactions and quaternary structure of protein complexes, as well as denatured TDMS to provide detailed proteoform information. Given the large number of AMPK combinations expected *in vivo* due to its diverse isoforms, nucleotide binding sites, and phosphorylation patterns, native TDMS offers a promising approach to elucidate the full complexity of this essential regulatory system. The ability to resolve kinase proteoform-ligand complexes opens new avenues for studying kinase regulation and function. Beyond AMPK, many protein kinases also undergo dynamic structural and proteoform changes that are challenging to capture using conventional approaches. We envision the TDMS-based strategy can serve as a powerful platform to elucidate a wide range of kinase complexes.

## Methods

### Materials and Reagents

All chemicals and reagents, including ammonium acetate (AA), formic acid (FA), and chloroform (CHCl_3_), were purchased from MilliporeSigma (Burlington, MA, USA) unless otherwise noted. Acetonitrile (ACN), isopropanol (IPA), and methanol (MeOH) were purchased from Fisher Scientific (Fair Lawn, NJ). Aqueous solutions were made in nanopore deionized water from Milli-Q water (MilliporeSigma). Micro Bio-Spin P-30 columns were purchased from Bio-Rad (Hercules, CA, USA).

### Cell Culturing, Protein Expression, and Affinity Purification

The recombinant αβγ complex was expressed from the tricistronic plasmid, pET28 MBP NAAEF-AMPKa1 (13-476,529-550)-b2(76-272)-g1(24-327), which was a gift from Karsten Melcher (Addgene plasmid # 177850; http://n2t.net/addgene:177850 ; RRID:Addgene_177850).^18^ Cultures were streaked on LB agar plates with kanamycin and incubated at 37°C overnight. Individual colonies were picked and added to LB liquid media (50 μg/mL kanamycin) and grown at 37°C until visually turbid. Plasmids were then purified using QIAprep Spin Miniprep Kit (Qiagen, Hilden, Germany).

The plasmid was transformed into One Shot BL21 (DE3) Competent *E. coli* (Thermo Fisher, Waltham, MA, USA). Cells were streaked on LB agar plates with kanamycin and incubated at 37°C overnight. Individual colonies were picked and added to 5 mL LB liquid media and grown at 37°C until visually turbid. The liquid culture was added to another 955 mL LB liquid media. Flasks were incubated at 37°C with agitation, monitoring growth until an OD_600_ of 0.8-1.0 was achieved. Cultures were then incubated on ice for 10 min, and 100 μL of 1 M isopropyl β-D-1-thiogalactopyranoside was added. Cultures were then incubated at 18°C with agitation overnight. Cultures were harvested, and cells were lysed using cell lysis buffer (20 mM Tris-HCl, 150 mM NaCl, pH 7), pelleted and then frozen at -80°C for additional lysis.

For purification of MBP-tagged AMPK, cell pellets were resuspended in lysis buffer (10 mM Tris-HCl, 150 mM NaCl, 5 mM MgCl_2_, 1 mM EDTA, 10% (v/v) glycerol) and sonicated. The homogenized lysate was then cleared by centrifugation for 40 min at 20,000 x*g* and filtered through a syringe filter (1.0 μm pore size). Filtered lysate was then added to a gravity column loaded with amylose resin (New England Biolabs, Ipswich, MA, USA), and MBP-tagged AMPK was eluted using the elution buffer (10 mM Tris-HCl, 150 mM NaCl, 5 mM MgCl_2_, 1 mM EDTA, 10% (v/v) glycerol, 40 mM maltose). The purified products were concentrated and stored at -80°C for further MS analysis.

### Native Top-Down MS Analysis

Native protein samples were prepared by buffer-exchanging into 300 mM AA solution using Bio-Spin columns. Samples were analyzed by nanoelectrospray ionization (nanoESI) via direct infusion using a TriVersa Nanomate system (Advio BioSciences, Ithaca, NY, USA) coupled to a solariX XR 12T Fourier transform ion cyclotron resonance mass spectrometer (FTICR-MS, Bruker Daltonics, Billerica, MA, USA)^44,69^. For the nanoESI source, the desolvating gas pressure was set at 0.6 PSI and the voltage was set to 1.65 kV. The source dry gas flow rate was set to 4 L/min at 120 °C. For the source optics, the capillary exit, detector plate, funnel 1, skimmer voltage, funnel RF amplitude, octopole frequency, octopole RF amplitude, collision cell RF frequency, and collision cell RF amplitude were optimized at 190 V, 200 V, 150 V, 120 V, 300 Vpp, 2 MHz, 600 Vpp, 1.4 MHz, and 2000 Vpp, respectively. Mass spectra were acquired with the mass range of 200-15000 *m/z* or 200-8000 *m/z*. For complex-up analysis, quadrupole was set at 5000 *m/z* to cut off low-molecular weight species. For complex-down analysis, an isolation window of 40 *m/z* was used for selecting the precursor ions. For CAD MS/MS experiments, an energy from 40 to 60 V was set to generate fragment ions. For ECD MS/MS experiments, the ECD pulse length, bias, and lens were set to 0.03 s, 1.5-3.5 V, 15-35 V, respectively. Acquisition size varied from 512k to 2M-words of data.

### Denatured Top-Down MS Analysis

Denatured protein samples were first pelleted from stock solution using chloroform/methanol/water precipitation. Protein pellets were reconstituted in 75% H_2_O/10% ACN/10% IPA/5% FA to make a 1 µg/µL solution. An Impact II quadrupole time-of-flight mass spectrometer (Q-TOF, Bruker Daltonics) coupled with a nanoACQUITY ultra-performance liquid chromatography (UPLC) system (Waters Corporation, Milford, MA, USA) was used for online LC-MS/MS analysis. Samples were first diluted with H_2_O to 0.2 µg/µL, and 200 µg of protein was injected into a home-packed PLRP (PLRP-S, Agilent Technologies) reversed-phase column (150 mm length × 250 µm I.D., 10 µm particle size, 1000 Å pore size). The mobile phases were 0.2% FA in H_2_O (A) and 0.2% FA in ACN (B) using a gradient of 10-10-70-95-95-10-10 % B in 0-2-24-24-27-28-30 minutes at a flow rate of 12 μL/min. Mass spectra were acquired at a scan rate of 1.0 Hz over 300-3000 *m/z*. For the ESI source, the end plate offset, capillary, nebulizer, dry gas, and dry temperature were set at 500 V, 4500 V, 0.5 bar, 4.0 L/min, and 200 °C, respectively. For tune settings, the funnel 1 RF, isCID energy, funnel 2 RF, hexapole RF, quadrupole ion energy, collision energy, collision RF, transfer time, and pre-pulse storage were set at 300 Vpp, 10 eV, 300 Vpp, 5 eV, 10 eV, 2600 Vpp, 120 µs, and 20 µs, respectively. For targeted CAD MS/MS experiments of AMPKα, an 80 *m/z* isolation window was applied to select the 7 topmost abundant charge states. Collision energies were set to values ranging from 15 to 25 eV.

A solariX XR 12T FTICR mass spectrometer coupled to a TriVersa Nanomate system was used for offline denatured top-down MS/MS experiments. For the nanoESI source, the desolvating gas pressure was set at 0.45 PSI and the voltage was set to 1.5-1.55 kV. The source dry gas flow rate was set to 3 L/min at 180 °C. For the source optics, the capillary exit, detector plate, funnel 1, skimmer voltage, funnel RF amplitude, octopole frequency, octopole RF amplitude, collision cell RF frequency, and collision cell RF amplitude were optimized at 240 V, 220 V, 150 V, 50 V, 300 Vpp, 5 MHz, 600 Vpp, 2 MHz, and 2000 Vpp, respectively. Mass spectra were acquired with an acquisition size of 2M in the mass range of 200-4000 *m/z*. An isolation window of at least 4 *m/z* was used for selecting the precursor ion. For CAD MS/MS experiments, an energy from 12 to 35 V was set to generate fragment ions. For ECD MS/MS experiments, the parameters for ECD pulse length, ECD bias, and ECD lens were set to 0.01-0.03 s, 0.5-1.5 V, 5-15 V, respectively.

### Data Analysis

Data were processed and analyzed using Compass DataAnalysis v. 4.3 and MASH Native v. 1.1.^70^ Maximum Entropy algorithm was used for spectra deconvolution with corresponding resolution (Q-TOF: 60,000; FTICR: 4,500 - 200,000, depending on acquisition size). FTMS algorithm (signal-to-noise ratio (S/N) threshold: 4; relative intensity threshold: 0.01%; absolute intensity threshold: 100) was applied to determine the most abundant mass of detected ions. The abundance of each proteoform was determined using DataAnalysis. MS/MS data were analyzed using MASH Native for proteoform sequencing and PTM localization. Fragments were identified using eTHRASH algorithm with a mass tolerance of 20 ppm. All fragments were manually validated.

## Acknowledgments

This work is supported by the National Institute of Health (NIH) NIH R01 GM117058 (to both Y.G. and S.J.). Y.G. would also like to acknowledge NIH/NHLBI R01 HL109810 and S10 OD018475. B.K. was supported by the European Union grant 101068151, Top-AMPK, HORIZON-MSCA-2021-PF-01. E.A.C. would like to acknowledge support from the NIH Chemistry-Biology Interface Training Program NIH T32GM152341. H.T.R. would like to acknowledge support from the National Heart, Lung, and Blood Institute of the NIH under Award Number T32HL007936 through the UW-Madison Cardiovascular Research Center. C.U. acknowledges funding through EU Horizon 2020 ERC StG-2017 759661 grant.

## Data Availability

The mass spectrometry proteomics data generated in this study have been deposited to the ProteomeXchange Consortium via the PRIDE partner repository with the data set identifier PXD062177 and MassIVE repository with identifier MSV000097430.

## Contributions

H.-J.C., B.K., and Y.G. designed research; H.-J.C. and L.J.B. performed MS analysis; B.K. and L.J.B. expressed and purified proteins; H.-J.C. and L.J.B. analyzed data; E.A.C., H.T.R., M.S.F., D.S.R., Z.G., and M.-D.W. provided input on experiments and evaluated results; J.W. and C. U. provided critical comments of the manuscript; H.-J.C., S.J., and Y.G. wrote the paper with comments from other co-authors.

