## Supporting Information for "Structural Heterogeneity of Proteoform-Ligand Complexes in AMP-Activated Protein Kinase Uncovered by Integrated Top-Down Mass Spectrometry"

### Supplementary Tables

**Table S1. AMPK proteoform-ligand complexes identified in native TDMS analysis.** Experimental most abundant mass (mean ± standard deviation, n = 3), calculated most abundant mass, and mass error for the identified complex. Abbreviations: methionine (Met); phosphorylation (P, phospho); adenosine monophosphate (AMP).

| Proteoform-Ligand Complex | Experimental Mass (Da) | Calculated Mass (Da) | ΔMass (Da) | Annotation | |
|---|---|---|---|---|---|
| | | | | PTMs | Ligands |
| AMPK αβγ | 153713 ± 1 | 153714 | <1 | Met removal*3 | |
| AMPK αβγ + P | 153793 ± 4 | 153794 | <1 | Met removal*3, phospho*1 | |
| AMPK αβγ + AMP | 154060 ± 4 | 154061 | 1 | Met removal*3 | AMP*1 |
| AMPK αβγ + AMP + P | 154139 ± 3 | 154141 | 1 | Met removal, phospho*1 | AMP*1 |
| AMPK αβγ + 2 AMP | 154407 ± 2 | 154408 | <1 | Met removal*3 | AMP*2 |
| AMPK αβγ + 2 AMP + P | 154486 ± 3 | 154488 | 2 | Met removal, phospho*1 | AMP*2 |

**Table S2. Dissociated AMPK subunit proteoforms and ligand binding identified in complex-up analysis.** Experimental most abundant mass, calculated most abundant mass, and mass error for the identified complex. Abbreviations: methionine (Met); phosphorylation (P, phospho); adenosine monophosphate (AMP).

| Subunits | Experimental Mass (Da) | Calculated Mass (Da) | ΔMass (Da) | Annotation | |
|---|---|---|---|---|---|
| | | | | PTM | Ligand |
| AMPK β | 22341.0 | 22341.4 | 0.5 | Met removal | |
| AMPK β + P | 22421.2 | 22421.4 | 0.2 | Met removal, phospho | |
| AMPK γ | 34659.9 | 34658.5 | 1.4 | Met removel | |
| AMPK αγ | 131376.2 | 131372.1 | 4.1 | α: Met removal γ: Met removal | AMP*1 |
| AMPK αγ + AMP | 131720.6 | 131719.2 | 1.4 | α: Met removal γ: Met removal | AMP*1 |
| AMPK αγ + 2 AMP | 132079.3 | 132066.2 | 13.1 | α: Met removal γ: Met removal | AMP*2 |
| AMPK αβ | 119052.5 | 119055.1 | 2.6 | α: Met removal β: Met removal | |
| AMPK αβ + P | 119132.4 | 119135.0 | 2.6 | α: Met removal β: Met removal, phospho | |
| AMPK αβ + AMP | 119406.6 | 119402.1 | 4.5 | α: Met removal β: Met removal | AMP*1 |
| AMPK αβ + AMP + P | 119488.4 | 119482.1 | 6.3 | α: Met removal β: Met removal, phospho | AMP*1 |

**Table S3. AMPK β and γ subunit proteoforms identified in complex-up analysis using in-source collisionally activated dissociation (IS-CAD).** Experimental most abundant mass, calculated most abundant mass, and mass error for the identified complex. Abbreviations: methionine (Met); phosphorylation (P, phospho); gluconoylation (G, glucono); phosphogluconoylation (PG, phosphoglucono); adenosine monophosphate (AMP).

| Subunit Precursor | Experimental Mass (Da) | Calculated Mass (Da) | ΔMass (Da) | Error (ppm) | Annotation | |
|---|---|---|---|---|---|---|
| | | | | | PTM | Ligand |
| AMPK β (12+) | 22341.6 | 22341.4 | 0.1 | 5.8 | Met removal | |
| AMPK β + P (12+) | 22421.6 | 22421.4 | 0.2 | 6.8 | Met removal, phospho | |
| AMPK γ (16+) | 34658.8 | 34658.5 | 0.3 | 8.5 | Met removal | |
| AMPK γ + G (16+) | 34835.9 | 34835.5 | 0.3 | 9.7 | Met removal, glucono | |
| AMPK γ + PG (16+) | 34915.8 | 34915.5 | 0.3 | 8.4 | Met removal, phosphoglucono | |
| AMPK γ + AMP (16+) | 35005.9 | 35005.5 | 0.3 | 9.1 | Met removal | AMP |

**Table S4. AMPK fragments identified in complex-down analysis.** For MS/MS spectra of complex-down analysis in **Figure S4** and **Figure S5**, fragment ion type, charge, experimental monoisotopic mass, calculated monoisotopic mass, and mass error were listed.

| Subunit (CE) | Ion | Charge | Exp Mass (Da) | Calc Mass (Da) | Error (ppm) |
|---|---|---|---|---|---|
| β, 12+ (CE=40V) | y42 | 3 | 4910.67 | 4910.67 | -0.6 |
| | y42 | 2 | 4910.66 | 4910.67 | 2.1 |
| | b27 | 2 | 3034.55 | 3034.54 | -1.9 |
| | b32 | 2 | 3598.94 | 3598.94 | 0.0 |
| | b36 | 2 | 4052.10 | 4052.10 | -0.3 |
| | b41 | 3 | 4595.44 | 4595.44 | 0.1 |
| | b42 | 3 | 4710.47 | 4710.47 | -0.5 |
| | b42 | 2 | 4710.47 | 4710.47 | 0.5 |
| | b43 | 3 | 4823.55 | 4823.56 | 0.3 |
| | b47 | 3 | 5235.75 | 5235.71 | -6.4 |
| | b55 | 3 | 6300.22 | 6300.22 | 0.0 |
| | b61 | 3 | 7022.58 | 7022.54 | -5.6 |
| | b61 | 4 | 7022.53 | 7022.54 | 1.4 |
| γ, 16+ (CE=50V) | y20 | 1 | 1981.18 | 1981.18 | 1.1 |
| | y23 | 1 | 2339.29 | 2339.29 | 0.9 |
| | y33 | 1 | 3456.92 | 3456.92 | -1.0 |
| | y41 | 2 | 4395.50 | 4395.47 | -6.4 |
| | y53 | 2 | 5781.16 | 5781.18 | 2.3 |
| | y68 | 3 | 7564.09 | 7564.09 | 0.0 |
| | y74 | 3 | 8284.39 | 8284.40 | 1.4 |
| | y82 | 3 | 9122.84 | 9122.89 | 6.0 |
| | y88 | 3 | 9876.27 | 9876.26 | -1.0 |
| | y142 | 6 | 15831.39 | 15831.40 | 1.1 |
| | b18 | 1 | 2076.88 | 2076.88 | 1.1 |
| | b20 | 1 | 2303.04 | 2303.05 | 1.5 |
| | b30 | 1 | 3376.60 | 3376.62 | 6.4 |
| | b30 | 2 | 3376.62 | 3376.62 | 0.1 |
| | b34 | 2 | 3805.85 | 3805.84 | -1.6 |
| | b35 | 2 | 3904.90 | 3904.91 | 3.5 |
| | b54 | 3 | 5990.05 | 5990.07 | 2.8 |
| | b102 | 5 | 11846.06 | 11846.10 | 4.0 |
| | b105 | 6 | 12208.32 | 12208.30 | -1.7 |
| | b114 | 6 | 13102.85 | 13102.76 | -6.3 |
| γ , 16+ (CE=60V) | y20 | 1 | 1981.18 | 1981.18 | 3.0 |
| | y23 | 1 | 2339.29 | 2339.29 | 0.2 |
| | y33 | 1 | 3456.93 | 3456.92 | -2.4 |
| | y68 | 3 | 7564.10 | 7564.09 | -0.6 |
| | y88 | 3 | 9876.15 | 9876.26 | 11.4 |
| | b18 | 1 | 2076.88 | 2076.88 | 1.0 |
| | b19 | 1 | 2189.95 | 2189.96 | 4.2 |
| | b20 | 1 | 2303.04 | 2303.05 | 1.6 |
| | b30 | 1 | 3376.62 | 3376.62 | 0.2 |

**Supplementary Figures**

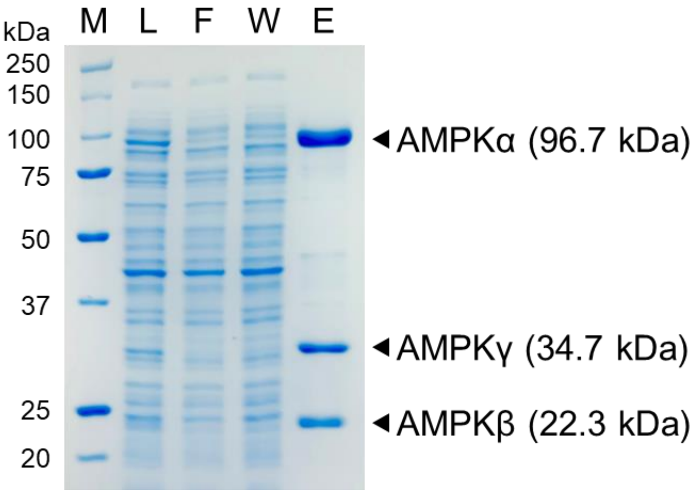

**Figure S1. SDS-PAGE analysis of AMPK affinity purification.** Samples loaded in each lane were labeled. M: marker, L: lysate, F: flow through, W: wash, E: elution.

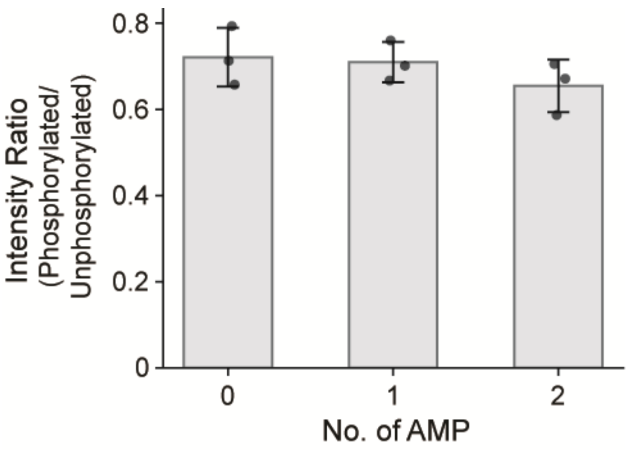

**Figure S2. Comparison of phosphorylation levels across different ligand binding states.** The intensity ratio of phosphorylated and unphosphorylated AMPK when binding to 0, 1, or 2 AMP molecules. Data are presented as mean ± standard deviation (n = 3).

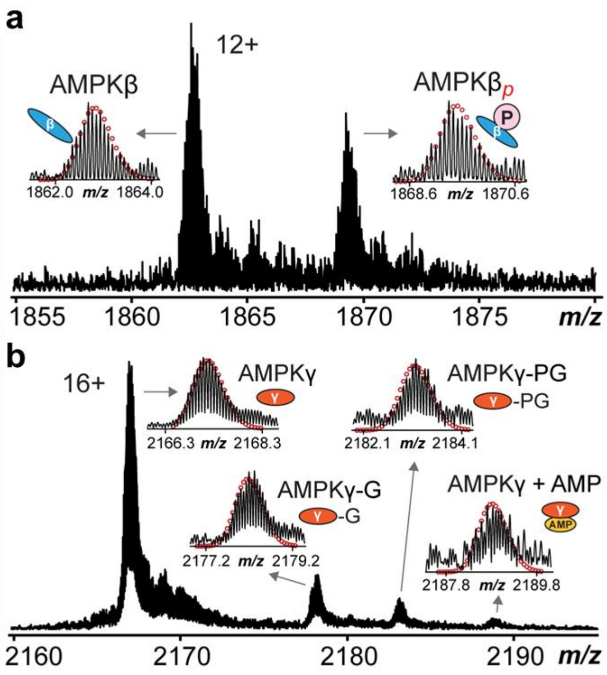

**Figure S3. Representative mass spectrum of complex-up analysis using in-source collisionally activated dissociation.** The mass spectrum was acquired with an acquisition size of 2M. Zoomed-in views of (a) AMPK β ($z$ = 12+) and (b) AMPK γ ($z$ = 16+). P: phosphorylation, G: gluconoylation, PG: phosphogluconoylation, AMP: adenosine monophosphate. Theoretical isotopic distributions are indicated by the red circles.

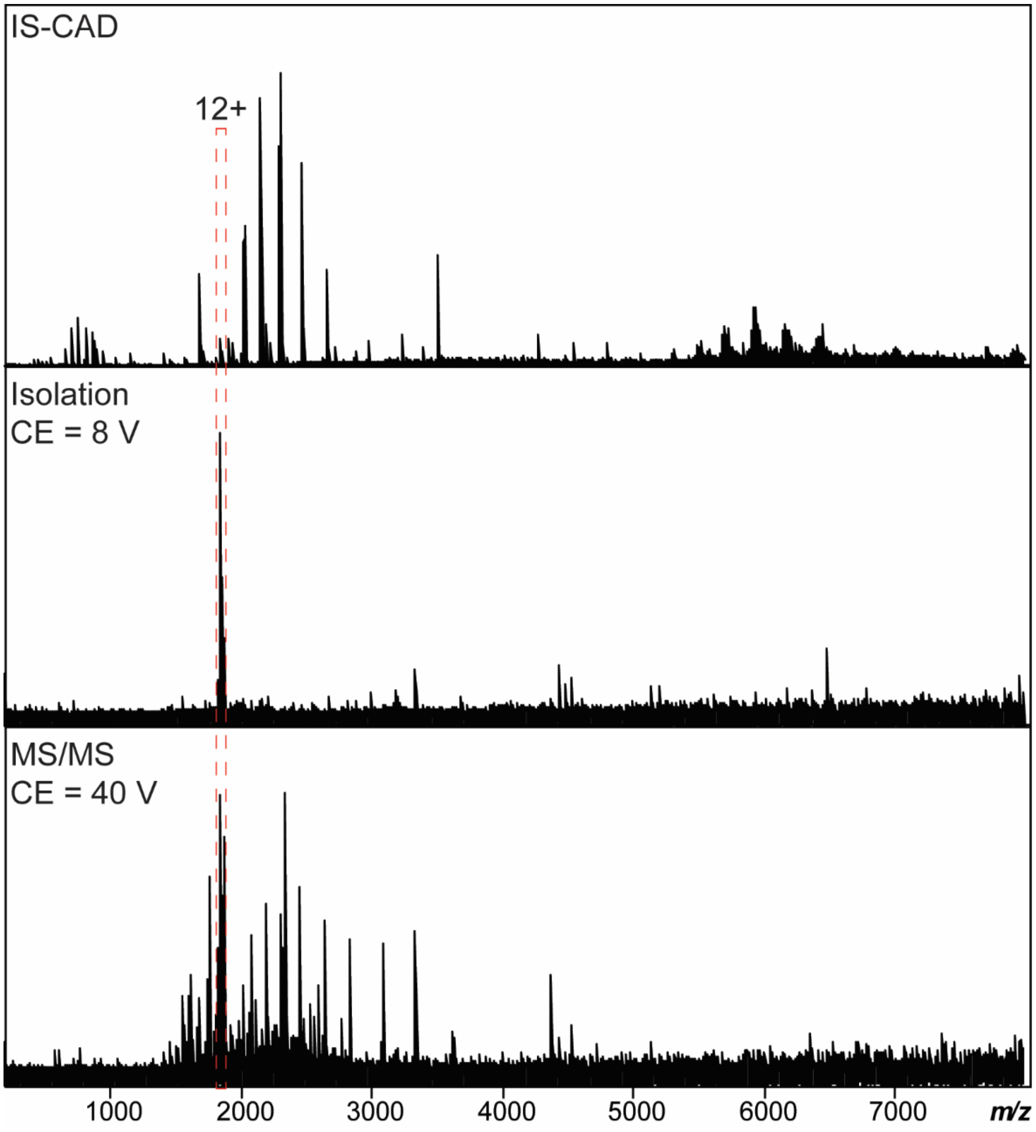

**Figure S4. IS-CAD and complex-down analysis of AMPK β subunit.** MS/MS characterization of the isolated β subunit ($z$ = 12+). Subunit dissociation was induced by applying IS-CAD in the funnel skimmer region (funnel 1 = 180 V and skimmer 1 = 160 V). For precursor isolation and complex-down MS/MS, a CAD energy of 8 V and 40 V were applied in the collision cell, respectively.

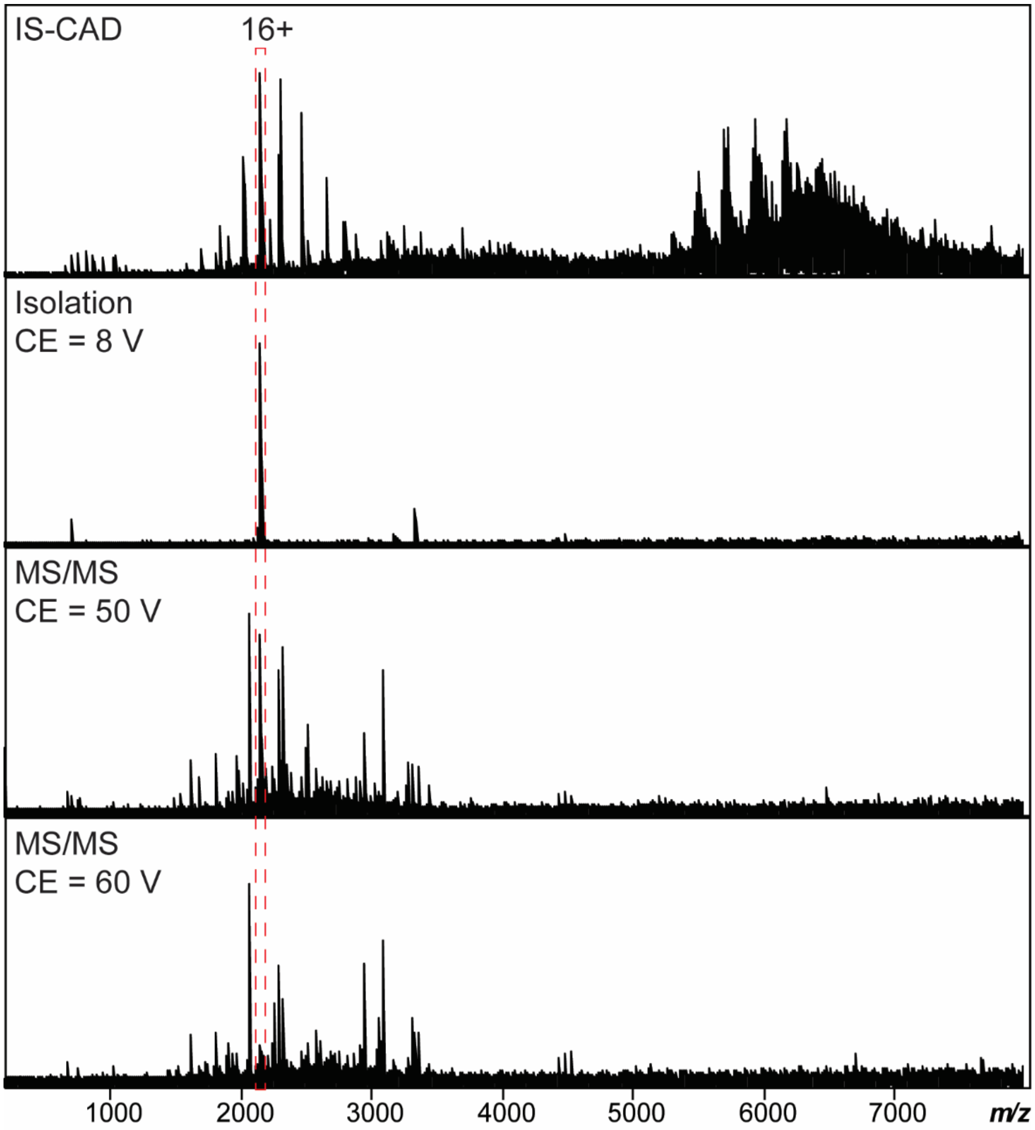

**Figure S5. IS-CAD and complex-down analysis of AMPK γ subunit.** MS/MS characterization of the isolated AMPK γ subunit ($z$ = 16+). Subunit dissociation was induced by applying IS-CAD in the funnel skimmer region (funnel 1 = 180 V and skimmer 1 = 170 V). For precursor isolation and complex-down MS/MS, a CAD energy of 8 V, 50 V, and 60 V were applied in the collision cell, respectively.

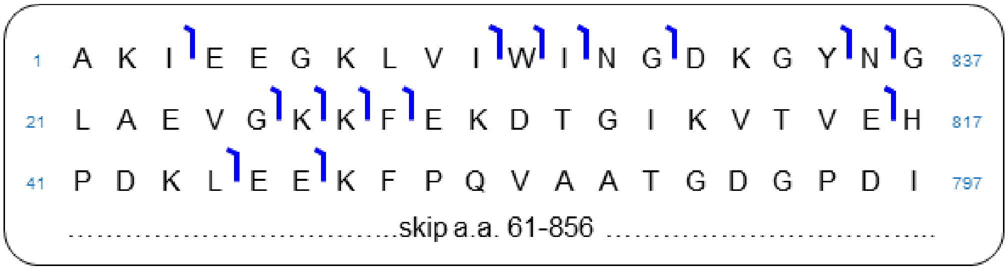

**Figure S6. Native top-down fragmentation map of AMPK α subunit.** Intact AMPK complex was fragmented using electron-capture dissociation (ECD) resulting in sequence informative *c* ions confirmed within a 20-ppm mass error tolerance. The sequence table shows the characterization of AMPK α subunit. The native top-down ECD data for AMPK α generated 14 *c* ions achieving 1.6% total bond cleavage.

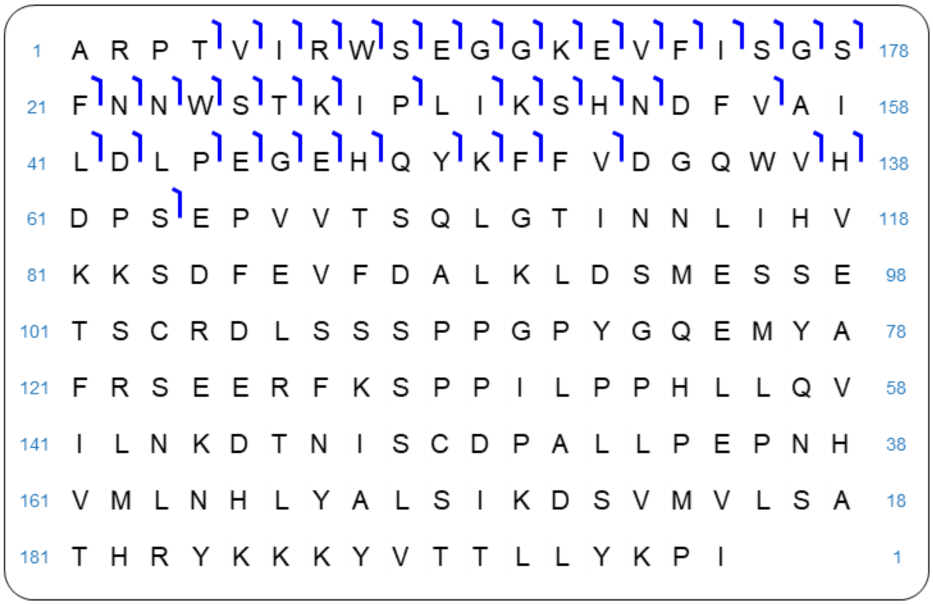

**Figure S7. Native top-down fragmentation map of AMPK β subunit.** Intact AMPK complex was fragmented using ECD resulting in sequence informative *c* ions confirmed within a 20-ppm mass error tolerance. The sequence table shows the characterization of AMPK β subunit. The native top-down ECD data for AMPK β generated 45 *c* ions achieving 23% total bond cleavage.

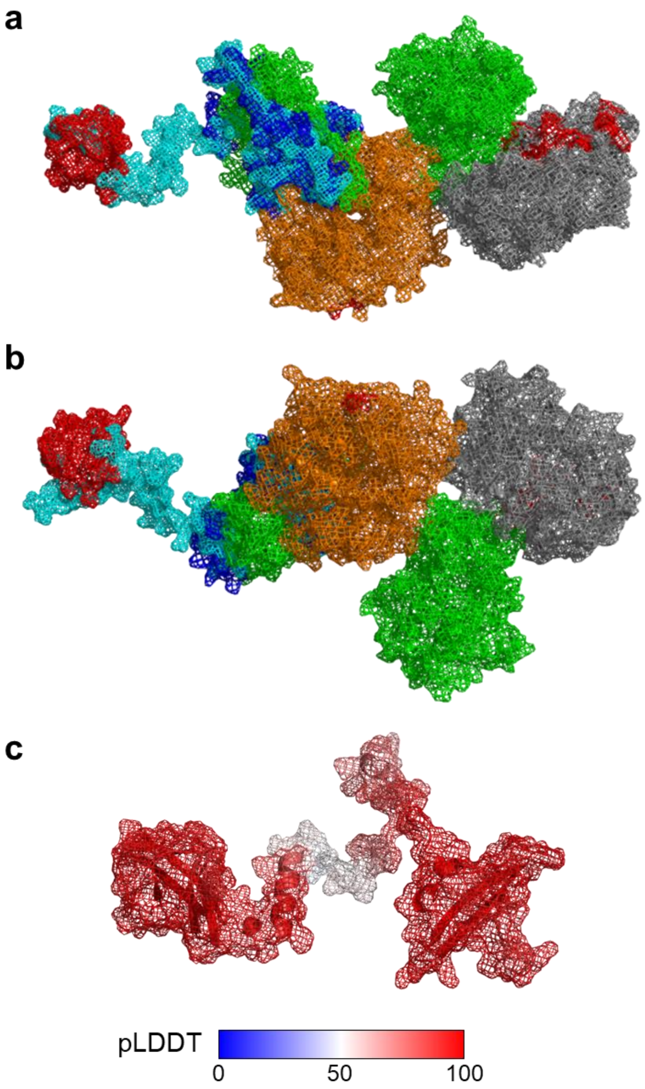

**Figure S8. Representative AMPK structure annotated with native top-down ECD fragmentation sites.** AMPK heterotrimeric complex (PDB: 7M74) was aligned and merged with full-length β subunit predicted by AlphaFold (AF-O43741-F1-v4). (a) Front view. (b) Back view. The experimental structure of α, β, and γ is labeled in green, dark blue, and orange, respectively. The maltose binding protein tag fused to the N-terminus of the α is labeled in gray. The predicted structure of the β subunit is labeled in cyan. Bond cleavage sites are labeled in red. (c) AlphaFold predicted β subunit (AF-O43741-F1-v4) colored by pLDDT values.

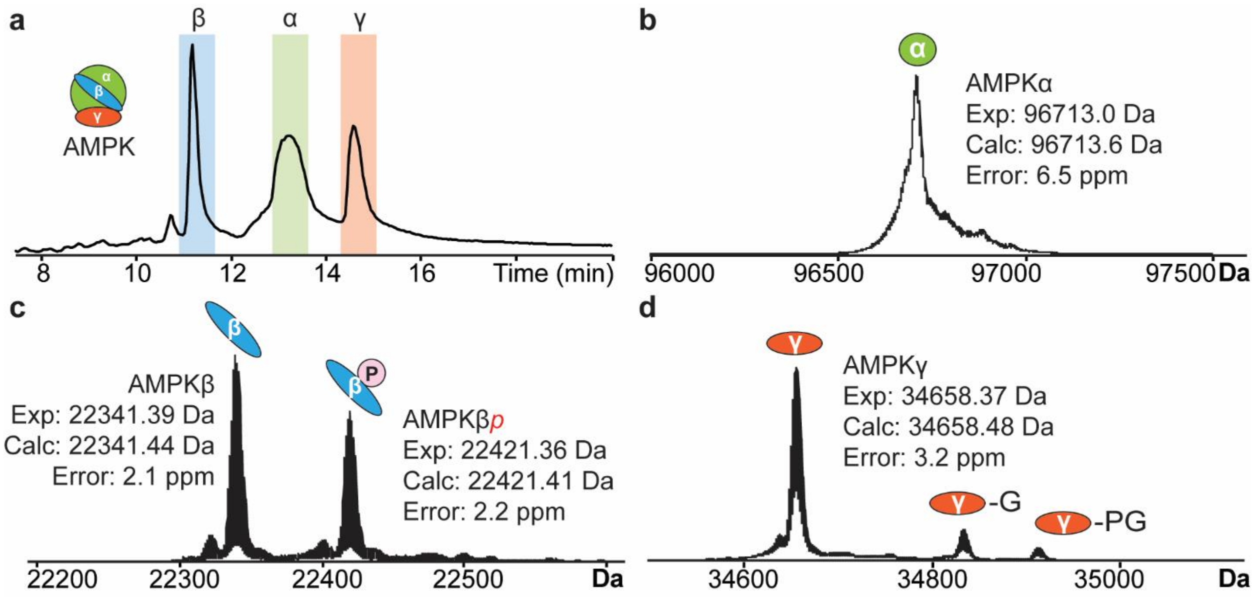

**Figure S9. Denatured TDMS analysis of AMPK subunits using online RPLC-Q-TOF MS.** (a) Total ion chromatogram of AMPK. The elution windows of the 3 subunits are labeled. Deconvoluted spectra of (b) α, (c) β, and (d) γ subunits. Experimental masses, calculated masses, and mass errors were reported.

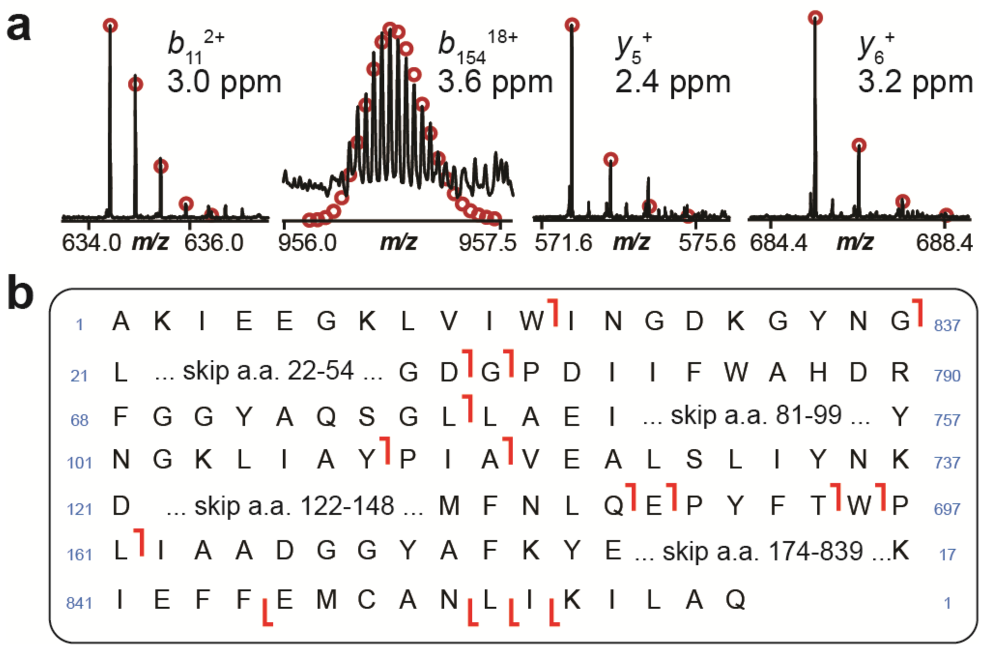

**Figure S10. Denatured TDMS analysis of AMPK α subunit using RPLC-Q-TOF MS.** (a) Representative CAD fragment ions ($b_{11}^{2+}$, $b_{154}^{18+}$, $y_5^{+}$, $y_6^{+}$) from denatured TDMS analysis. The isotopic fitting is shown with red circles and mass errors are reported. (b) Sequence map of the α subunit annotated with identified CAD fragments.

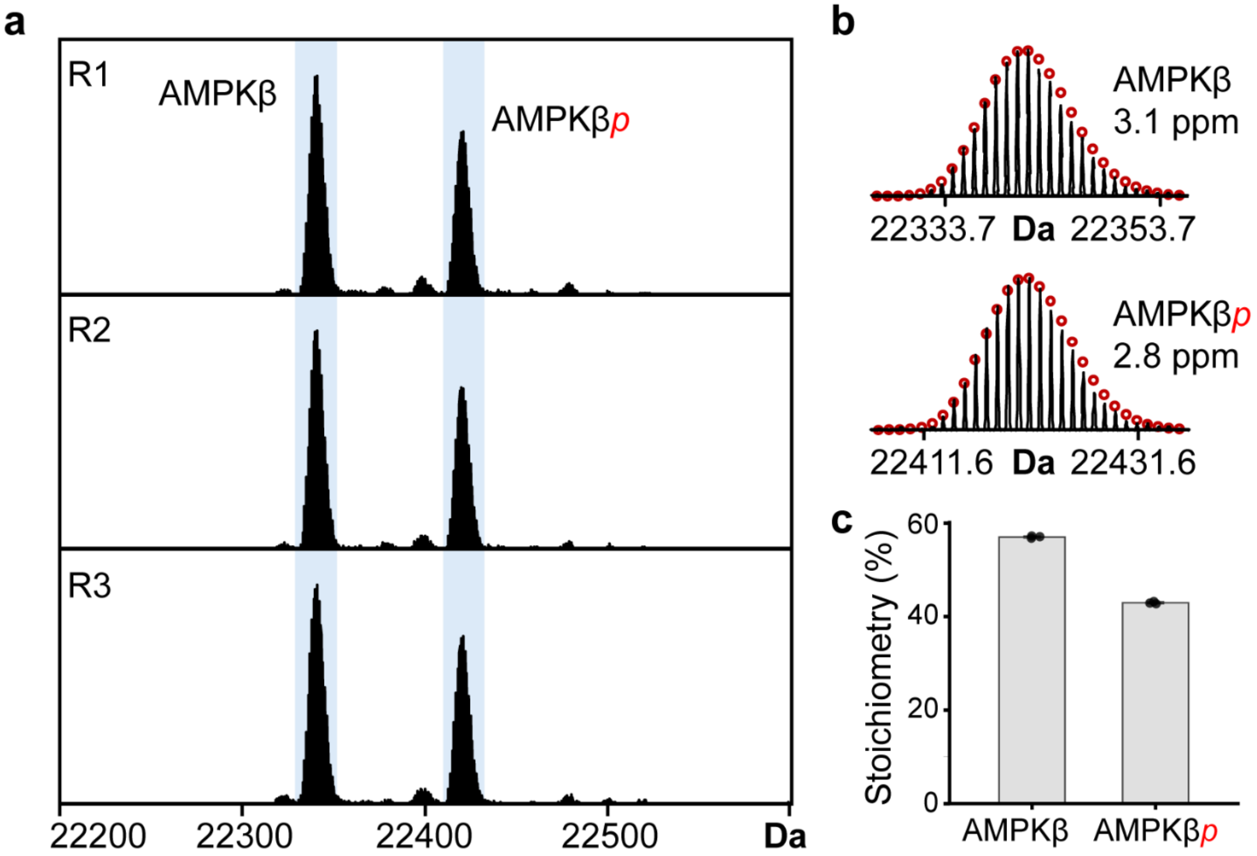

**Figure S11. Proteoform stoichiometry of AMPK β subunit.** (a) Representative deconvoluted mass spectra of AMPK β proteoforms (R1-3: technical triplicates). AMPKβ: unphosphorylated, AMPKβ*p*: monophosphorylated. (b) Isotope distribution of identified AMPK β proteoforms overlaid with the theoretical isotopic fitting (red circles). Mass errors are reported. (c) Proteoform stoichiometry of unphosphorylated and monophosphorylated AMPK β subunit. Data are presented as mean ± standard deviation (n = 3).

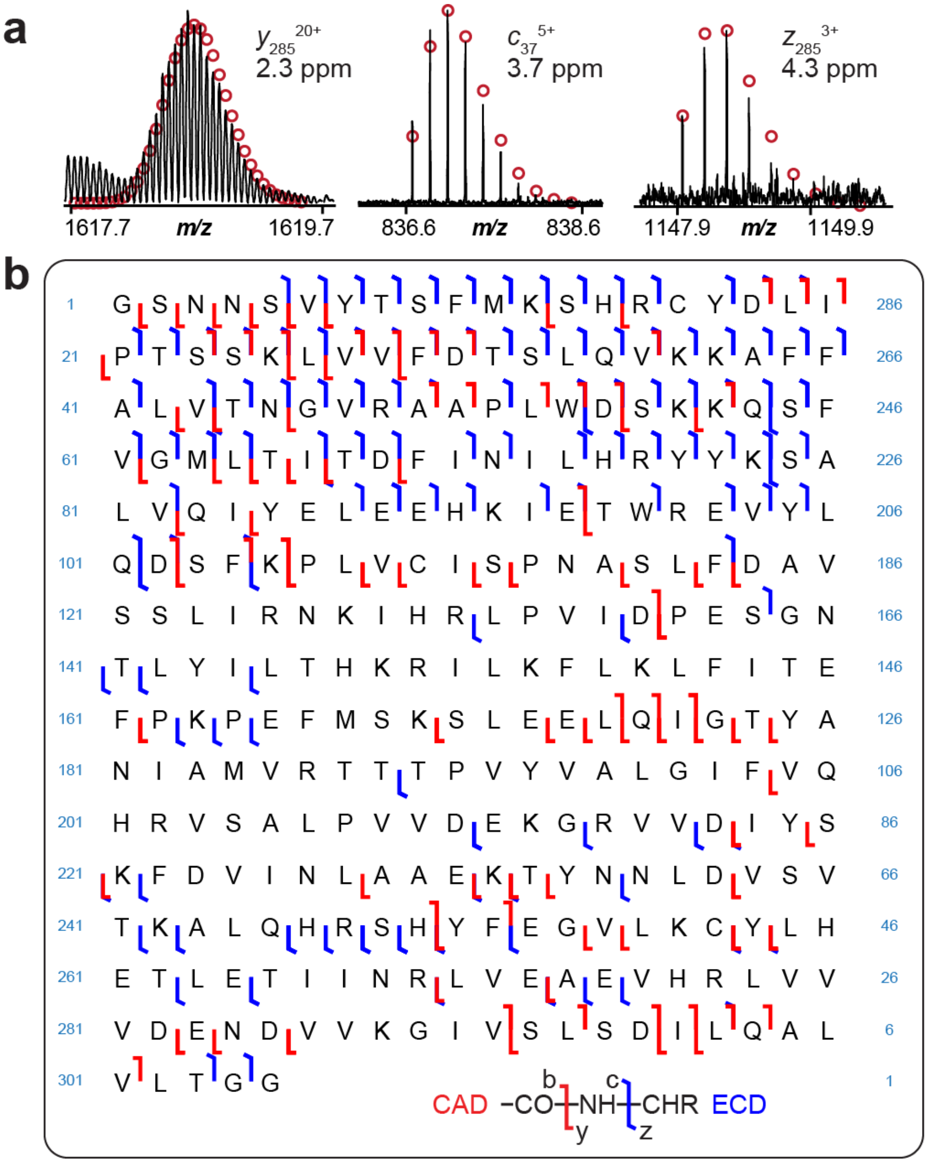

**Figure S12. Denatured TDMS analysis of AMPK γ subunit using FTICR-MS/MS.** (a) Representative CAD and ECD fragment ions ($b_{285}^{20+}$, $c_{37}^{5+}$, $z_{33}^{3+}$) from denatured TDMS analysis. The isotopic fittings are shown in red circles and mass errors are reported. (b) Sequence map of the γ subunit annotated with identified CAD and ECD fragments.
